# Oral Lisinopril Raises Tissue Levels of ACE2, the SARS-CoV-2 Receptor, in Healthy Male and Female Mice

**DOI:** 10.1101/2021.10.19.465025

**Authors:** Steven D. Brooks, Rachel L. Smith, Aline da Silva Moreira, Hans C. Ackerman

**Affiliations:** Physiology Unit, Laboratory of Malaria and Vector Research, National Institute of Allergy and Infectious Diseases, Rockville, Maryland, USA

**Keywords:** Angiotensin Converting Enzyme 2, Angiotensin Converting Enzyme inhibitor, Angiotensin Receptor Blocker, COVID-19, SARS-CoV-2

## Abstract

Angiotensin-converting enzyme 2 (ACE2) is the established cellular receptor for SARS-CoV-2. However, it is unclear whether ACE1 inhibitors (e.g., lisinopril) or angiotensin receptor blockers (e.g., losartan) alter tissue ACE2 expression. This study sought to determine whether lisinopril or losartan, as monotherapies or in combination, change tissue levels of ACE2 in healthy male and female mice.

Mice received lisinopril (10 mg/kg/day), losartan (10 mg/kg/day), or both for 21 days via drinking water. A control group received water without drug. ACE2 protein index (ACE2 protein / total protein) was determined on small intestine, lung, kidney, and brain. Oral lisinopril increased ACE2 protein index across all tissues (p < 0.0001 vs control). In contrast, the combination of lisinopril plus losartan did not increase ACE2 levels in any tissue (p = 0.89 vs control) and even decreased tissue expression of the *Ace2* gene (p < 0.001 vs control). Tissue ACE2 remained elevated in mice 21 days after cessation of lisinopril (p = 0.02). Across both cohorts, plasma ACE2 did not correlate with ACE2 protein index in any tissue. A sex difference was observed: kidney ACE2 levels were higher in males than females (p < 0.0001).

Oral lisinopril increases ACE2, the cellular receptor for SARS-CoV-2, in tissues that are relevant to the transmission and pathogenesis of COVID-19. Remarkably, the addition of losartan prevented lisinopril-induced increases in ACE2 across tissues. These results suggest that ACE inhibitors and angiotensin receptor blockers interact to determine tissue levels of ACE2.

## 1. INTRODUCTION

Angiotensin-converting enzyme 2 (ACE2) is an established receptor and entry point for both SARS-CoV-1 and the novel SARS-CoV-2 coronaviruses.^1–3^ The spike proteins on the viral envelope bind the ACE2 receptor, and the virus replicates efficiently in cells expressing ACE2.^1^ Human tissue histological profiling reveals ACE2 to be highly expressed on lung alveolar epithelial cells and on enterocytes of the small intestine, as well as on arterial and venous endothelium.^4^ SARS-CoV-2 can enter vascular endothelium in engineered human blood vessel organoids and human kidney organoids via ACE2.^5^ SARS-CoV-2 is also associated with endothelial inflammation,^6,7^ which may give rise to the clinical findings of thromboembolism^8^ and disseminated intravascular coagulation.^9^

Given the widespread abundance of ACE2 in tissue epithelial and endothelial cells, and the role of ACE2 as the entry site for SARS-CoV-2, there has been much speculation regarding whether ACE inhibitors and/or angiotensin receptor blockers (ARB) may alter ACE2 tissue abundance and thereby change the risk of transmission or development of severe complications.^10–12^ Recent clinical studies of patients with COVID-19 have not identified a clear relationship between ACE inhibitor use or ARB use and disease risk or severity,^13–15^ and current guidelines support continuance of ACE inhibitors or ARB during infection.^16,17^ The design of human trials and the development of clinical guidelines regarding ACE inhibitor and ARB use have been limited by the lack of preclinical data on how ACE inhibitors and ARB change tissue abundance of ACE2.^18–20^ Therefore, the question of how these drugs may impact tissue expression and abundance of ACE2 remains of fundamental interest.

The primary goal of this study was to determine whether lisinopril, an oral ACE inhibitor, or losartan, an oral ARB, changes the tissue abundance of ACE2, and whether these changes resolve after cessation of the drug. The tissues studied were the lung and small intestine, which have been identified as portals of entry for SARS-CoV-2 and sites of primary disease pathogenesis;^21–23^ the kidney, selected for its role in the angiotensin pathway^24^ and because renal failure is a complication of severe COVID-19;^25^ and the brain, due to the neurological symptoms and sequalae identified during acute and long-haul COVID-19.^26^ The secondary goals of this study were to determine whether tissue ACE2 levels differ between tissues, whether plasma ACE2 correlates with tissue ACE2, and whether tissue ACE2 levels differ between male and female mice. The findings we present below demonstrate that lisinopril raised ACE2 levels in tissues when given alone, but not when given in combination with losartan. ACE2 levels varied substantially between tissues, and plasma ACE2 did not correlate with tissue ACE2. We found kidney ACE2 levels to be greater in males than in females. Together, these results provide controlled experimental data demonstrating the impact of ACE inhibition and angiotensin receptor blockade on tissue ACE2 expression in mice and highlights a potential benefit of ACE inhibitor/ARB combination therapy in the setting of a SARS-CoV-2 pandemic.

## 2. METHODS

### 2.1 Use of laboratory mice

The protocols used in this study were performed in accordance with the National Institutes of Health guide for the care and use of Laboratory animals (NIH Publications No. 8023, revised 1978). All experiments and protocols using laboratory mice were reviewed and approved by the NIAID Division of Intramural Research Animal Care and Use Committee (DIR ACUC).

### 2.2 Experimental design

The experiment utilized a factorial design: five male and five female mice comprised each drug treatment group (lisinopril, losartan, lisinopril and losartan combined, or vehicle) at each time point (Day 21 or Day 42). These forty male and forty female eight-week old C57Bl/6J mice (Jackson Laboratory) were fed standard chow and treated for 21 days with either drinking water containing lisinopril (10 mg/kg/day; Exelan Pharmaceuticals), drinking water containing losartan (10 mg/kg/day; Aurobindo Pharma), combination (10 mg/kg/day of each drug), or standard drinking water (vehicle control). On day 21, forty animals were euthanized for collection of plasma and tissues, while the others transitioned to standard drinking water for an additional 21 days to assess whether drug-induced changes in ACE2 resolve after drug cessation.

A pilot study of five male and five female eight-week-old C57Bl/6J mice was performed to measure average daily drinking water intake. This value was used to calculate the initial drug concentration needed to achieve a 10mg/kg/day dosage. Each week of drug treatment during the main study, every mouse was weighed (Supplemental Fig. 2) and water consumption was measured on a per-cage basis (Supplemental Fig. 3). This information was used to adjust the drug concentration weekly to maintain consistent dosing throughout the course of the study.

### 2.3 Tissue collection and processing

Each mouse was euthanized via bilateral thoracotomy while under anesthesia with 4% isoflurane. 1 mL of blood was drawn into an EDTA tube via cardiac puncture of the right ventricle. 25 mL of cold phosphate-buffered saline was administered via transcardial perfusion to remove blood from tissues prior to collection. Small intestine, lung, kidney, and brain were collected and flash-frozen for protein extraction, stored in 10% formalin for histological examination, or stored in RNAlater for gene expression studies. Plasma was separated from whole blood by centrifugation and frozen.

### 2.4 Measurement of ACE2 protein index

Flash-frozen lung, small intestine, kidney, and brain were homogenized at 4°C (Precellys Cryolys Evolution, Bertin Instruments) in lysis buffer (RIPA buffer, 1X, Cell Signaling). Tissue total protein concentration was measured by BCA assay (Pierce BCA). ACE2 tissue abundance was measured by ELISA (Abcam). To minimize the effects of inter-assay variation, all biospecimens from a given experimental day (21 or 42) were analyzed together on a single ELISA plate and BCA plate. ACE2 protein index was calculated by dividing the ELISA-measured concentration by the total protein concentration of each specimen. ACE2 concentration in plasma (pg/mL) was measured by ELISA (Abcam).

### 2.5 Measurement of Ace2 gene expression

mRNA was extracted (RNeasy, Qiagen) from small intestine, lung, kidney, and brain tissue sections, stored in RNAlater and converted to cDNA (SuperScript VILO IV, Invitrogen). Expression of *Ace2* and the reference gene *Gapdh* were measured by Reverse Transcriptase droplet digital PCR (Prime ddPCR assays, QX200, Bio-Rad). Gene expression was quantified as the transcript ratio of *Ace2* to *Gapdh*.

### 2.6 Immunohistochemistry of tissue sections

Small intestine, lung, kidney, and brain tissue sections were fixed in 10% Formalin, embedded into paraffin, sliced into 6-8 μm sections by microtome, and stained with immunohistochemical antibodies for ACE2 (Sino Biological, 1:1000) to determine tissue prevalence and distribution.

### 2.7 Measurement of plasma renin activity

Plasma renin activity was measured using a fluorometric Renin Assay Kit (Abcam). Plasma renin activity was measured via cleavage of a fluorogenic substrate over 60 minutes at 37 degrees C and reported as renin concentration equivalent (ng/mL) using a reference renin standard provided by Abcam. Fluorescence measurements (excitation/emission = 540/590) were made by microplate reader (MD Gemini XPS).

### 2.8 Statistical Analyses

Data were tested for normality using Shapiro-Wilk test. Tissue ACE2 data were transformed by Box-Cox method before analysis. Multivariable analysis of variance was used to assess the effect of treatment on tissue ACE2 protein index, with tissue type and sex as covariates. Each drug treatment group was compared against vehicle control using Tukey post-hoc tests and adjusted p-values were reported.

The *Ace2*/*Gapdh* transcript ratio data were normal by Shapiro-Wilk test. Multivariable analysis of variance was used to assess the effect of treatment on the *Ace2*/*Gapdh* transcript ratio, with tissue type and sex as covariates. Each drug treatment group was compared against vehicle control using Tukey post-hoc tests and adjusted p-values were reported.

Body weights and water consumption were measured weekly. The effect of treatment and time on body weight and water consumption was assessed separately in male and female mice by Repeated Measures two-way ANOVA. The effect of treatment and sex on plasma renin activity was assessed separately at Day 21 and Day 42 by two-way ANOVA. Linear regression was used to measure the relationship between plasma ACE2 and ACE2 protein index in each tissue; covariates were sex and treatment.

A posthoc multivariate model, prompted by visual inspection of the data, was used to test the effect of sex on kidney ACE2 protein index across both cohorts and all treatment groups.2.9 *Supplemental Materials and Methods:* Detailed methodology and a comprehensive description of antibodies, PCR reaction components, and laboratory equipment and consumables are available in Appendix A: Supplemental Materials and Methods.

## 3. RESULTS

### 3.1. Tissue ACE2 protein index and *Ace2* gene expression differed by tissue type

To assess tissue-specific ACE2 abundance, ACE2 protein index was analyzed in the small intestine, kidney, lung, and brain of male and female vehicle-treated mice at day 21. ACE2 protein index differed significantly by tissue (p < 0.0001, two-way ANOVA); it was highest in the small intestine, followed by kidney, lung, and brain (Fig. 1A; Supplemental Table 1). To assess tissue-specific *Ace2* gene expression, the *Ace2*/*Gapdh* transcript ratio was analyzed in the small intestine, kidney, lung, and brain of male and female vehicle-treated mice at day 21. The *Ace2*/*Gapdh* transcript ratio differed significantly by tissue (p < 0.0001, two-way ANOVA); it was highest in the small intestine, similar in the kidney and lung, and lowest in the brain (Fig. 1B; Supplemental Table 2).

**Figure 1:**
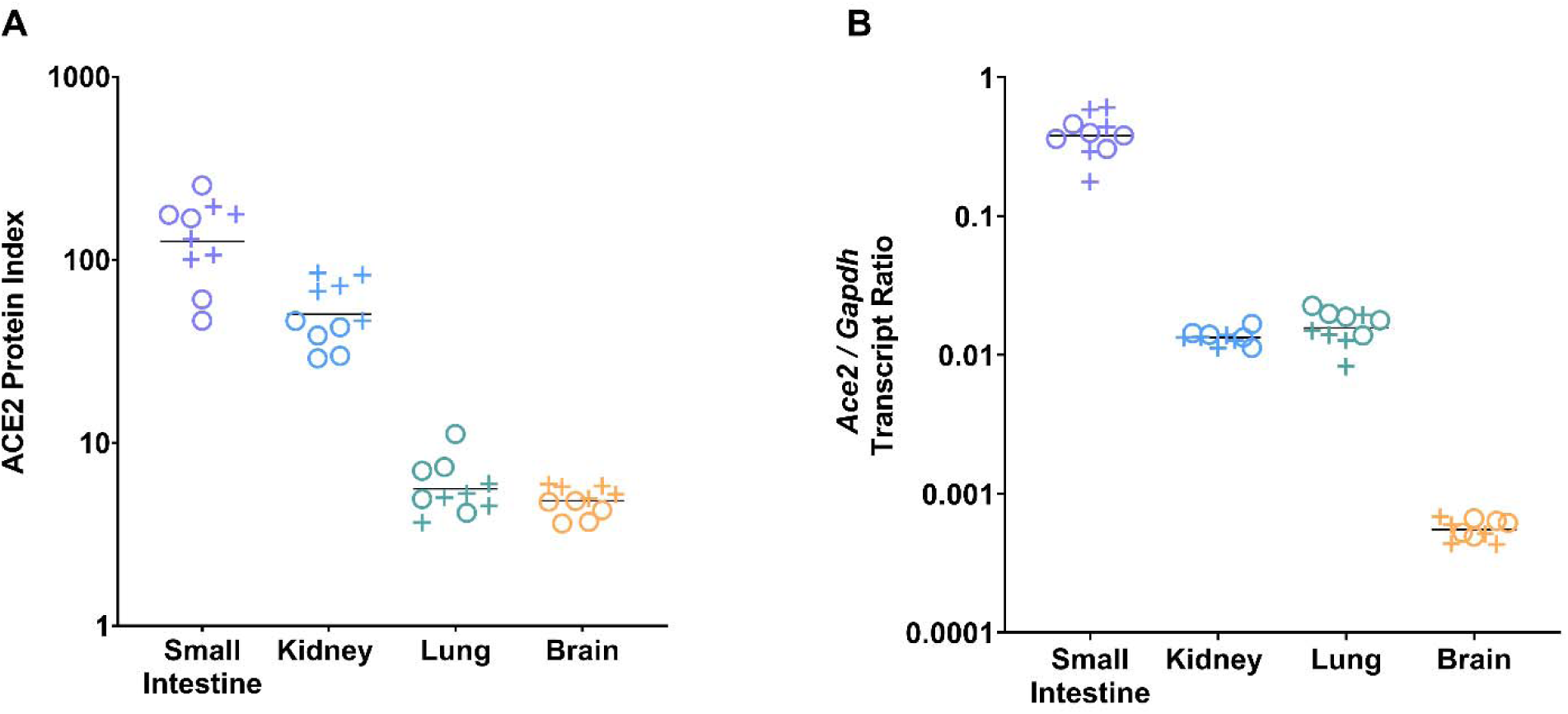
ACE2 protein index and *Ace2* gene expression differed by tissue type. **(A)** ACE2 protein level was measured by ELISA and normalized to BCA-determined total protein concentration to generate tissue-specific ACE2 protein indices for the small intestine, kidney, lung, and brain of vehicle-treated male (+) and female (o) animals (5 males, 5 females per group). Tissue ACE2 protein index values were multiplied by 10^6^ for display purposes. ACE2 protein index differed by tissue (p < 0.0001 by two-way ANOVA). ACE2 protein index (mean ± SD) was highest in the small intestine (1.41×10^−4^ ± 0.65 ×10^−4^), followed by kidney (5.40×10^−5^ ± 2.10×10^−5^), lung (5.91×10^−6^ ± 2.19×10^−6^), and brain (4.90×10^−6^ ± 0.83×10^−6^) (p_adj_ < 0.0001 for small intestine versus kidney, lung or brain by Tukey post-hoc test; p_adj_ = 0.02 for kidney versus lung and p_adj_ = 0.02 for kidney vs brain; no difference between lung and brain). (B) *Ace2* gene expression was measured by ddPCR and normalized to *Gapdh* and presented as the *Ace2*/*Gapdh* transcript ratio for the small intestine, kidney, lung, and brain of vehicle-treated male (+) and female (o) animals. *Ace2* expression differed by tissue (p < 0.0001 by two-way ANOVA). *Ace2* expression (mean ± SD) was highest in the small intestine (0.400 ± 0.131) compared to kidney (0.013 ± 0.002), lung (0.016 ± 0.004) or brain (0.001 ± 0.0001) (p_adj_ < 0.0001 by Tukey test).

Given the abundance of ACE2 in tissues from vehicle-treated mice, we preserved tissue segments in 10% buffered formalin and then sectioned and stained for ACE2 (Supplemental Fig. 1). In the small intestine, ACE2 was detected along the microvilli of the small intestine in sections from vehicle-treated control and lisinopril-treated mice. In the kidney, ACE2 was detected along the lumen of renal tubules, consistent with tubular epithelium. In the lung, ACE2 was detected in the alveolar epithelium. In the brain, ACE2 localized to the vasculature.

### 3.2 Lisinopril treatment raised ACE2 protein index in tissues, but the combination of lisinopril and losartan did not

ACE2 protein index was determined in the small intestine, kidney, lung, and brain of male and female mice after 21 days of treatment with lisinopril, losartan, combination, or vehicle (Fig. 2 and Supplemental Table 1). To test the effect of treatment on tissue ACE2 protein index, multivariate analysis of variance was performed (Table 1). Treatment affected ACE2 protein index (p < 0.0001). Lisinopril treated mice had higher ACE2 protein indices compared to mice treated with vehicle (p_adj_ < 0.0001 by Tukey post-hoc test). Losartan treated mice had a non-significant increase in ACE2 compared to vehicle controls (p_adj_ = 0.058). In contrast, the combination of lisinopril plus losartan did not raise tissue levels of ACE2 (p_adj_ = 0.89 vs vehicle control). Treatment had no effect on body weight (Supplemental Fig. 2), water consumption (Supplemental Fig. 3), or plasma renin activity (Supplemental Fig. 4).

**Figure 2:**
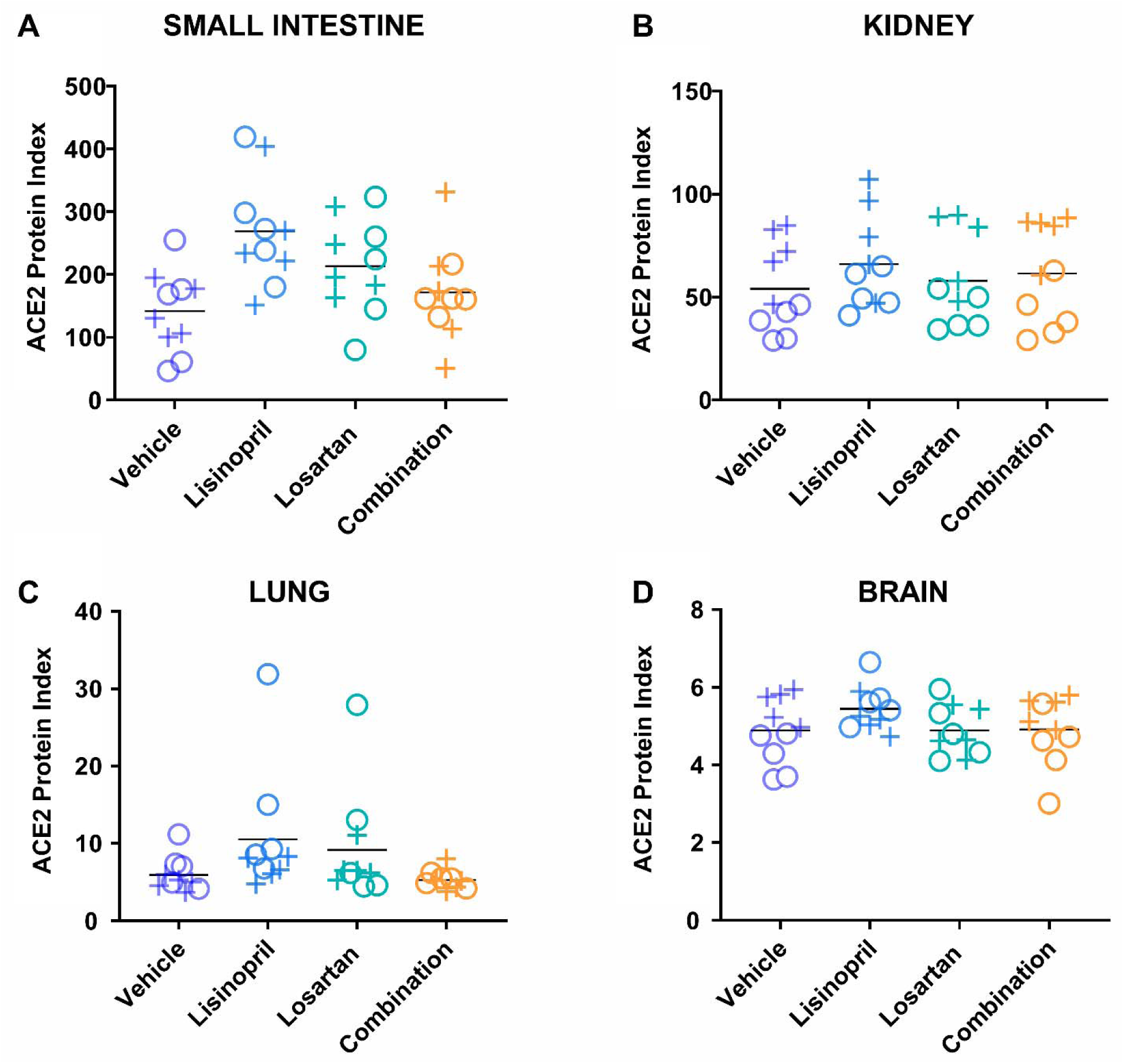
Lisinopril treatment raised the ACE2 protein index, but the combination of lisinopril and losartan did not. ACE2 protein index was measured in the small intestine (A), kidney (B), lung (C), and brain (D) of male (+) and female (o) mice after 21 days of treatment (5 males, 5 females per group). ACE2 protein index was different across treatment groups (p < 0.0001). Lisinopril-treated mice had higher ACE2 protein indices compared to mice treated with vehicle control (p_adj_ < 0.0001 by Tukey post-hoc test), while mice treated with the combination of lisinopril plus losartan were not different from mice treated with vehicle control (p_adj_ = 0.89). Tissue ACE2 protein index values were multiplied by 10^6^ for display purposes.

**Table 1.**
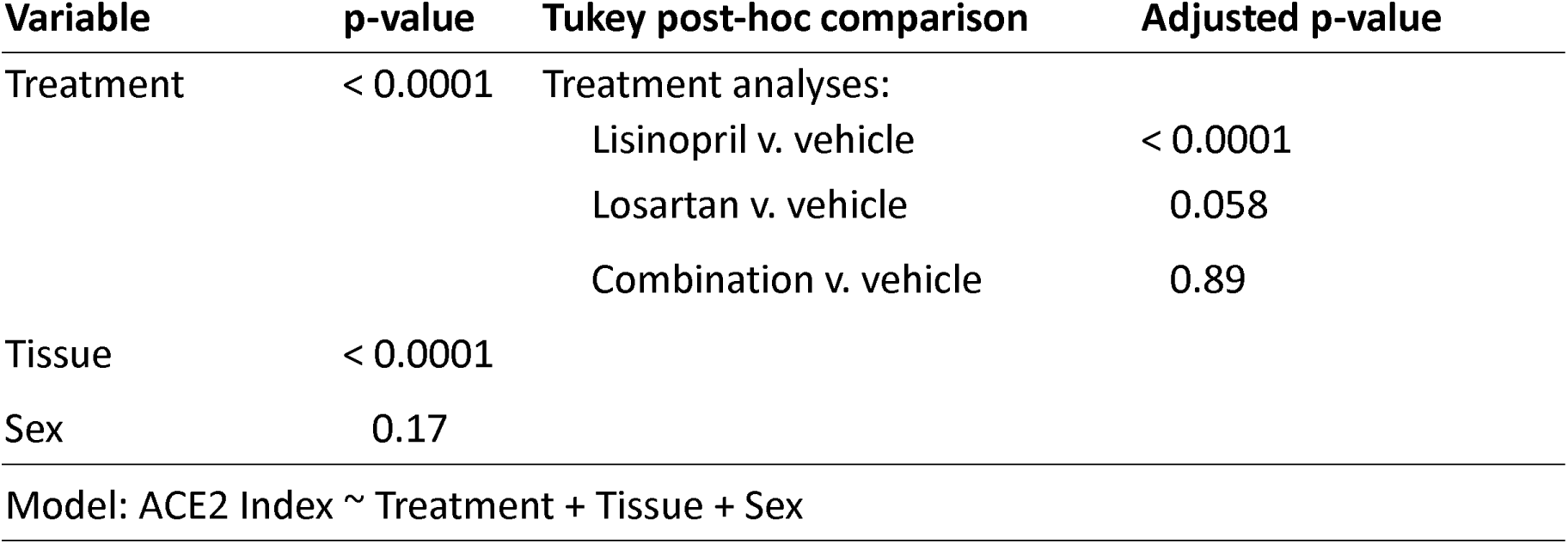
Effect of drug treatment on tissue ACE2 protein index on Day 21. **Lisinopril increased ACE2 protein index after 21 days of treatment**. After 21 days of treatment, ACE2 protein index was measured in small intestine, kidney, lung, and brain. The effect of treatment on ACE2 protein index across tissues was determined by ANOVA, which included terms for treatment group, tissue type, and sex in the model.

### 3.3 Lisinopril and losartan combination treatment suppressed *Ace2* gene expression in tissue

To further explore the different effects of lisinopril versus lisinopril/losartan combination on ACE2, we examined *Ace2* gene expression in the small intestine, kidney, lung, and brain at Day 21 (Fig. 3 and Supplemental Table 2). *Ace2* expression was highest in the small intestine (p_adj_ < 0.0001 against any other tissue. To assess the effect of treatment on tissue *Ace2* gene expression, a multivariate analysis of variance was performed that included terms for tissue type and sex (Table 2). Treatment affected *Ace2* gene expression (p = 0.001); tissue type was a significant factor (p < 0.0001), but sex was not (p = 0.733). The primary treatment effect was the lowering of *Ace2* expression by combination therapy compared to vehicle control (p_adj_ < 0.001) or lisinopril monotherapy (p_adj_ = 0.047). Neither lisinopril (p_adj_ = 0.60 by Tukey’s post-hoc test) nor losartan (p_adj_ = 0.104) monotherapy changed *Ace2* expression compared to vehicle.

**Figure 3:**
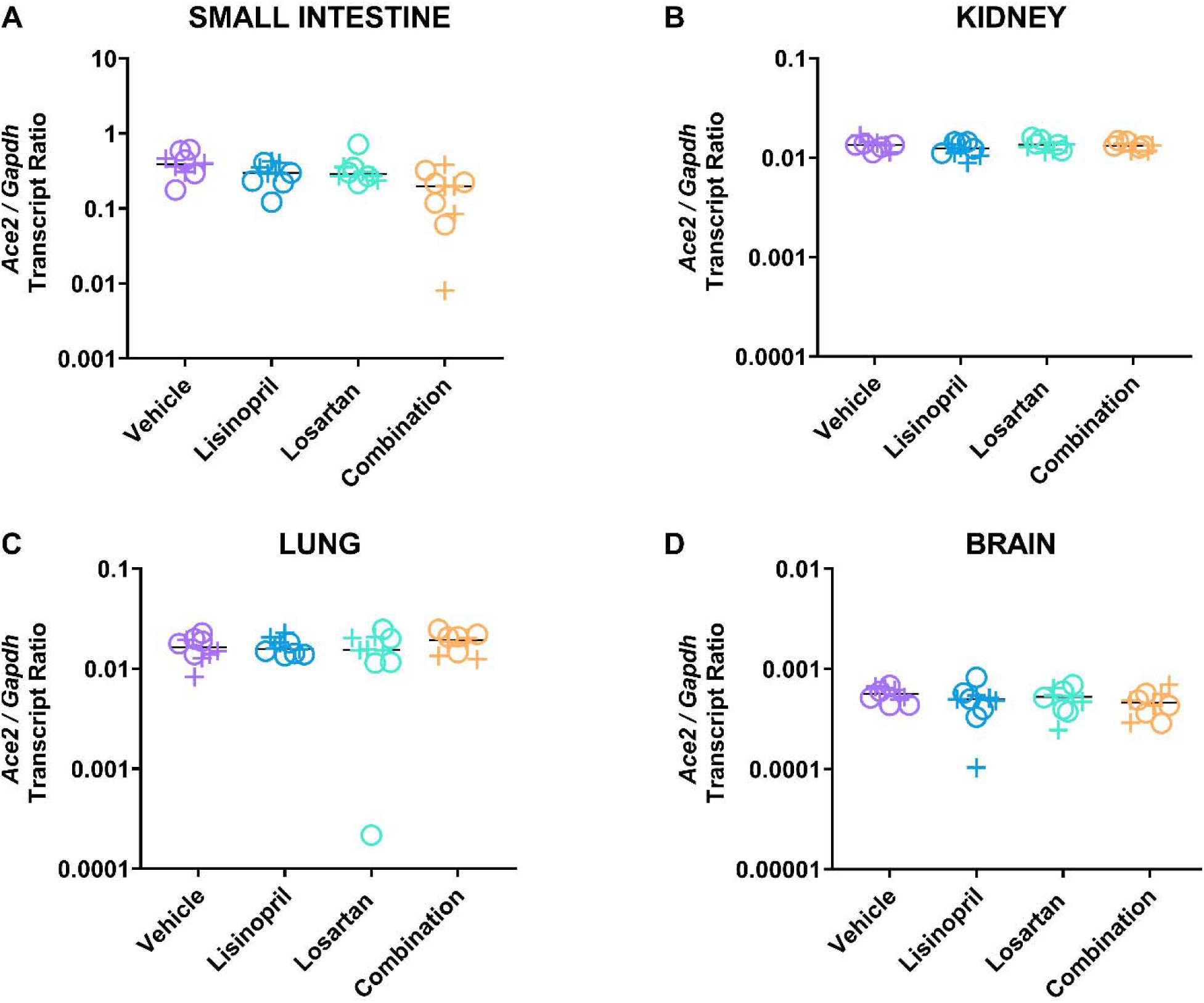
Lisinopril and losartan combination treatment suppressed *Ace2* gene expression. *Ace2* and *Gapdh* expression was measured in the small intestine of male (+) and female (o) mice by reverse transcriptase droplet digital PCR in the small intestine (A), kidney (B), lung (C), and brain (D) (5 males, 5 females per group). There was an effect of treatment (p = 0.001) and tissue (p < 0.0001), but no effect of sex (p = 0.733), on *Ace2*/*Gapdh* expression. Post-hoc testing showed no effect of lisinopril or losartan on gene expression compared to vehicle. Combination-treated mice had a significantly lower *Ace2*/*Gapdh* transcript ratio compared to vehicle (p_adj_ < 0.001) or lisinopril monotherapy (p_adj_ = 0.047) by Tukey post-hoc test. The effect was largest in the small intestine, which had the highest levels of expression (p_adj_ < 0.0001 versus any tissue by Tukey Test).

**Table 2.** Effect of drug treatment on *Ace2* expression on Day 21. **Combination therapy reduced *Ace2* gene expression after 21 days of treatment**.After 21 days of treatment, *Ace2* expression (*Ace2*/*Gadph* expression ratio) was measured in small intestine, kidney, lung, and brain. The effect of treatment on the *Ace2*/*Gapdh* expression ratio across tissues was determined by ANOVA, which included treatment group, tissue type, and sex as terms in the model.

| Variable | p-value | Tukey post-hoc comparison | Adjusted p-value |
| --- | --- | --- | --- |
| Treatment | 0.001 | Treatment analyses:<br>Lisinopril v. Vehicle<br>Losartan v. Vehicle<br>Combination v. Vehicle<br>Combination v. Lisinopril | 0.600<br>0.104<br>< 0.001<br>0.047 |
| Tissue | < 0.0001 | Tissue analyses:<br>Sm. Intestine. v. Brain<br>Sm. Intestine. v. Kidney<br>Sm. Intestine. v. Lung | < 0.0001<br>< 0.0001<br>< 0.0001 |
| Sex | 0.733 |  |  |
| Model: ACE2 Index ~ Treatment + Tissue + Sex |  |  |  |

### 3.4 Drug-induced elevation of ACE2 protein index persisted 21 days after discontinuation of drug

To determine whether the treatment-induced changes in ACE2 protein index were reversible, we measured tissue ACE2 protein index 21 days after cessation of drug treatment (Fig. 4 and Supplemental Table 1). In multivariate analysis that included treatment group, tissue type, and sex (Table 3), prior treatment was associated with higher ACE2 levels (p = 0.013 by ANOVA). Specifically, mice previously treated with lisinopril or losartan had higher tissue ACE2 levels than mice previously treated with vehicle control (p_adj_ = 0.025, p_adj_ = 0.024, respectively, by Tukey post-hoc test). ACE2 levels in mice previously treated with the combination of lisinopril and losartan were not different from mice previously treated with vehicle control (p = 0.30).

**Figure 4:**
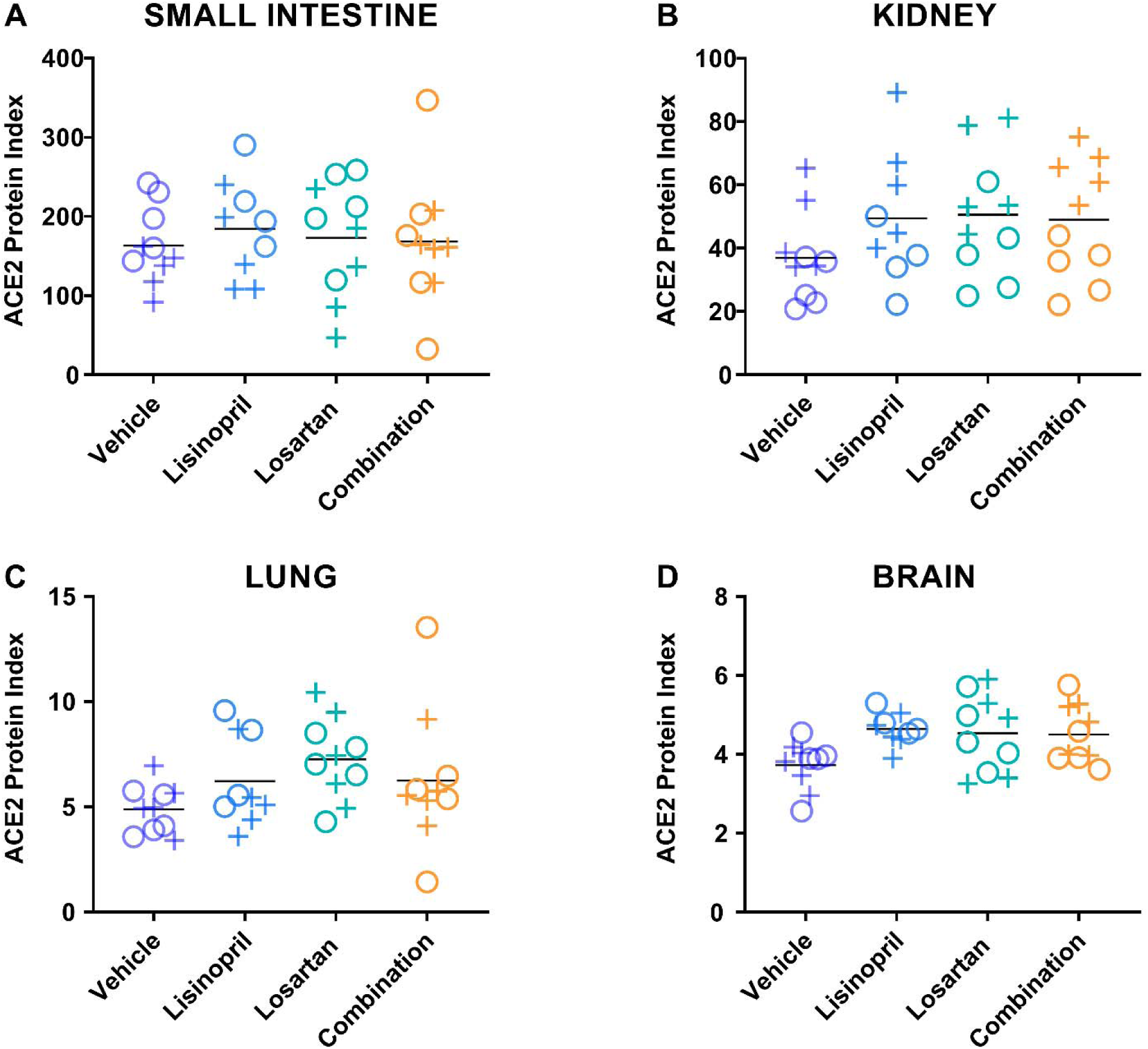
Drug-induced increases in ACE2 protein index persisted 21 days after discontinuation of drug treatment. ACE2 protein index was measured by ELISA in the (A) small intestine, (B) kidney, (C) lung, and (D) brain of male (+) and female (o) animals on day 42, 21 days after discontinuation of treatment with lisinopril, losartan, or lisinopril/losartan combination (n= 5 males, 5 females per group (except losartan females Day 42, n=4)). ACE2 protein index values were multiplied by 10^6^ for display purposes. The effect of treatment on ACE2 protein index remained significant 21 days after drug cessation (p = 0.013 by ANOVA). Specifically, mice previously treated with lisinopril (p_adj_ = 0.025) or losartan (p_adj_ = 0.024) had greater ACE2 levels than mice treated with vehicle control. Mice treated with the combination of lisinopril and losartan were not different from mice treated with vehicle control (p = 0.30).

**Table 3.**
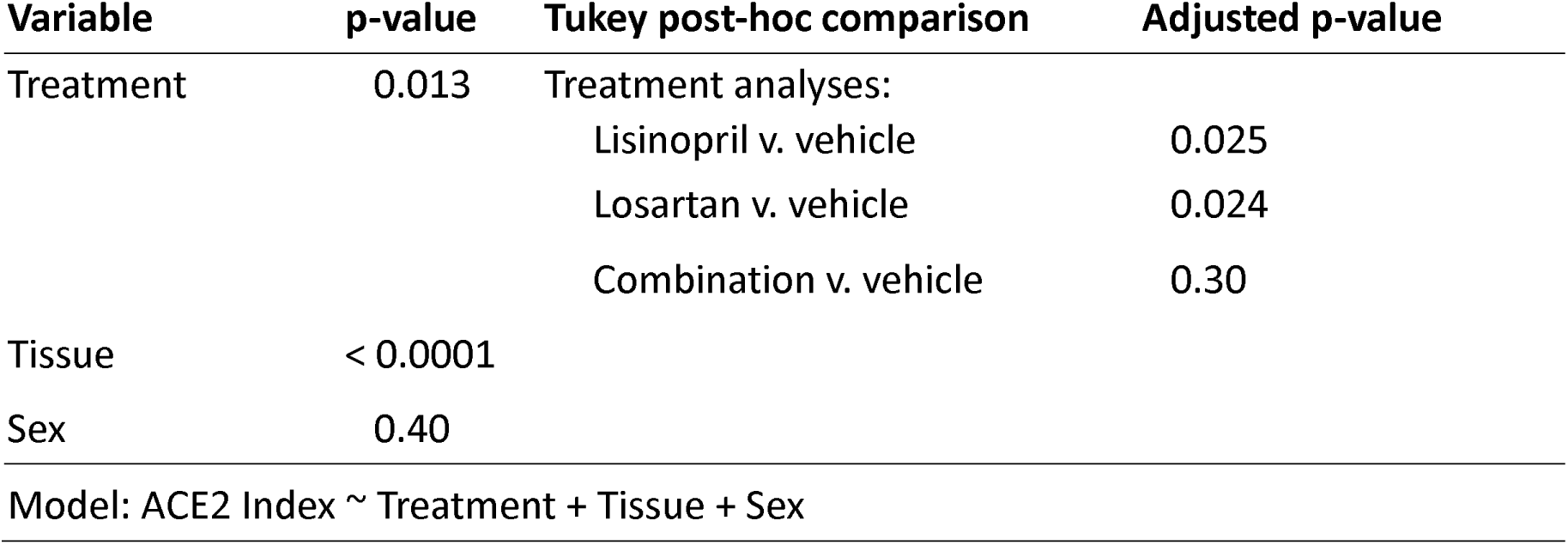
Effect of drug treatment on tissue ACE2 protein index on Day 42, 21 days after drug cessation. **Treatment-associated elevation of ACE2 protein index persisted 21 days after drug cessation**. After 21 days of treatment, drug treatment groups were switched to vehicle control. On day 42, ACE2 protein index was measured in small intestine, kidney, lung, and brain. The effect of treatment on ACE2 protein index across tissues was determined by ANOVA, which included treatment group, tissue type, and sex as terms in the model.

### 3.5 Plasma ACE2 did not correlate with tissue ACE2

Plasma ACE2 was measured in whole blood collected on Day 21 and Day 42 from male and female mice treated with lisinopril, losartan, combination, or vehicle. The relationships between plasma ACE2 and tissue ACE2 protein index in small intestine, lung, kidney, and brain, were analyzed by linear regression to identify whether plasma ACE2 could serve as a biomarker of tissue ACE2 (Supplemental Fig. 5). Linear regression analysis revealed that plasma ACE2 was not associated with tissue ACE2 in any tissue (small intestine p = 0.95; kidney p = 0.26; lung p = 0.90; brain p = 0.62).

### 3.6 Kidney ACE2 levels were higher in males versus females

Multivariate analysis of all treatment groups at day 21 or day 42 did not find sex to be a significant factor affecting tissue ACE2 levels (p = 0.17; p = 0.40). However, inspection of the data revealed a pattern of sex-based divergence in kidney ACE2 levels in both the day 21 and day 42 cohorts (data shown in Fig. 2 and Fig. 4). This prompted a post-hoc subgroup analysis focusing only on kidney ACE2 levels across both cohorts. Multivariate analysis of kidney ACE2 levels across both cohorts and all treatment groups revealed kidney ACE2 levels to be greater among males than females (p < 0.0001).

## 4. DISCUSSION

ACE2 serves as the cognate receptor for the SARS-CoV-2 spike protein on the apical surface of epithelial and endothelial tissues in humans. As a key cellular entry point for the virus, ACE2 is important for both transmission of virus from person to person and tissue-specific pathology caused by local viral entry. High levels of plasma ACE2 have been associated with increased risk of severe illness from COVID-19.^27^ There has been much speculation about how ACE inhibitors or ARB might alter expression of the ACE2 protein, and thereby potentially alter host susceptibility to infection with SARS-CoV-2 or the progression, severity, and tissue-specific pathology of COVID-19. However, few studies have directly measured the effect of ACE inhibition or angiotensin receptor blockade on ACE2 levels, especially outside the cardiovascular system.

To address the question of whether ACE inhibition and/or angiotensin receptor blockade alters tissue ACE2 expression, we measured tissue-specific changes in ACE2 abundance following treatment with an ACE inhibitor (lisinopril), an ARB (losartan), or the combination of both, compared to vehicle, in male and female mice. We found that 21 days of ACE inhibition with lisinopril increased tissue ACE2 expression compared to vehicle in analysis that included small intestine, kidney, lung, and brain. However, this increase in tissue ACE2 was prevented when losartan was given in combination with lisinopril. These treatment-related increases in tissue ACE2 were still detectable 21 days after discontinuation of the drugs.

A secondary objective of this study was to assess for sex differences in tissue ACE2 abundance and in response to drug treatment. When all tissues were examined together, sex was not significantly associated with tissue ACE2 levels; however, a tissue subgroup analysis revealed kidney ACE2 levels to be significantly higher in males compared to females among the drug-treated groups (p < 0.0001). Kidney ACE2 activity was previously reported to be greater in male versus female mice in the absence of drug treatment^28^, and a similar trend was observed in kidney tissue from human donors.^29^ Here we describe for the first time sex differences in kidney ACE2 in mice treated with ACE inhibitor and ARB.

Another secondary objective of this study was to test whether plasma ACE2 could serve as a biomarker for tissue ACE2 protein index. We tested for association between tissue ACE2 protein index and plasma ACE2 by linear regression but did not find a significant relationship between plasma ACE2 and ACE2 index in the small intestine, kidney, lung, or brain. Recently, clinical studies have used soluble ACE2 as a biomarker for activation of the Renin Angiotensin Aldosterone System (RAAS) or as a marker of tissue ACE2;^30,31^ however, our results in mice suggest that plasma ACE2 is not a suitable biomarker for tissue ACE2. While elevated plasma ACE2 has been associated with severe COVID-19 disease,^27^ our results did not find a link between plasma ACE2 and tissue ACE2 index in any tissue.

We measured renin activity in plasma from mice at Day 21 and Day 42 to test whether the lisinopril-induced increase in tissue ACE2 index could be related to hemodynamic effects of drug treatment on the renin-angiotensin system. We did not observe an increase in renin activity for any treatment group, suggesting that there were no hemodynamic changes induced by drug treatment. This observation was consistent with prior reports that ACE inhibitors and ARBs lower blood pressure in hypertensive mice but not in normotensive mice.^32–35^

Outside the cardiovascular system, we found that ACE2 was highly abundant around the lateral and apical margins of intestinal villi. Reports of fecal-oral transmission suggest that the intestinal tract may be a site of viral transmission,^36^ as well as a route of viral entry into epithelial cells leading to gastrointestinal symptoms.^37^ There is increasing interest in intestinal infection as a route of viral spread;^22^ among people taking ACE inhibitors, associated increases in small intestine ACE2 could potentially increase the risk of SARS-CoV-2 viral infection.

This study is the first to systemically evaluate the effect of ACE inhibition and angiotensin receptor blockade on ACE2 protein abundance in these four tissues along with plasma; moreover, we assessed for sex differences and evaluated whether drug-induced changes in tissue ACE2 resolve after drug cessation. Previously, pre-clinical studies examined the effect of ACE inhibitor and ARB treatment on *Ace2* gene expression in cardiac and lung tissue from rats and mice. The first found that in cardiac tissue, 12 days of lisinopril or losartan monotherapy increased *Ace2* gene expression, while the combination of lisinopril plus losartan did not.^38^ While we did not observe an increase in *Ace2* gene expression after monotherapy with lisinopril or losartan, we did observe a decrease in *Ace2* gene expression among mice treated with the combination of lisinopril and losartan compared to vehicle and lisinopril treatment, leading to similar differences in *Ace2* gene expression between treatment groups. Our finding that lisinopril increased tissue ACE2 protein index but combination therapy prevented the increase agree with the principal finding of the previous study, and further extends this finding to the protein level and across several relevant tissues.^38^ A second study reported that 21 days of either oral captopril (an ACE inhibitor) or oral candesartan (an ARB) up-regulated gene expression of *Ace2* and increased ACE2 enzymatic activity in lung tissue from healthy rats.^39^ In that study, combination therapy was not evaluated. Interestingly, they reported a sharp increase in *ACE2* gene expression in cultured human alveolar cells after 24 hours of exposure to captopril or candesartan that decreased to near-baseline by 48 hours despite the maintenance of elevated ACE2 protein.^39^ Another study found no changes in *Ace2* expression in lung, kidney, ileum or heart after 14 days of treatment with an ACE inhibitor,^40^ while a study in diabetic mice found eight weeks of ACE inhibitor treatment decreased *Ace2* mRNA in the diabetic kidney, but had no effect in the lung or heart on *Ace2* expression.^41^ Interestingly, that study also showed that diabetes increases ACE2 activity in kidney and lung.^41^ Our finding that elevated tissue ACE2 protein at day 21 in lisinopril-treated mice was not accompanied by a sustained increase in *Ace2* gene expression is consistent with these prior observations. Furthermore, we observed elevated ACE2 protein at Day 42 in the lisinopril- and losartan-treated groups compared to control even 21 days after cessation of drug. Taken together, the previous studies plus our present study indicate that ACE inhibitor or ARB monotherapy increases ACE2, as observed in rat tissue, mouse tissue, and cultured human cells, while combination therapy does not.

In contrast, another previous study in mice reported that an ACE inhibitor or ARB may actually reduce ACE2 in its membrane-bound form in the kidneys.^43^ This study examined the effects of two weeks of ACE inhibitor or ARB monotherapy on kidney and lung ACE2 gene expression, protein, and activity in tissue lysates from C57 mice.^43^ Similar to our results, no increase was found in *Ace2* mRNA in lung or kidney after ACE inhibitor or ARB treatment. In contrast, they found no difference in total kidney ACE2 from either drug, no change in lung ACE2 activity from either drug, but a significant decrease in membrane bound ACE2 along with an increase in cytosolic ACE2 in the kidney after either ACE inhibitor or ARB treatment. However, this study used Western blot (as opposed to quantitative ELISA) to estimate ACE2 protein quantity in kidney and lung, and did not normalize to BCA-quantified total protein^43^, making direct comparison to our study difficult. There was also no examination of drug withdrawal, and sex of study animals was not reported; whereas we observed significant sex differences on ACE2 protein index in the kidney.

While neither ACE inhibitors nor ARB bind directly to ACE2, they may modulate ACE2 expression indirectly by changing the circulating levels of angiotensin-II, the major substrate for ACE2. ACE inhibitors decrease circulating levels of angiotensin-II; in contrast, ARBs increase circulating levels of angiotensin-II.^18,42^ However, treatment with lisinopril, losartan, or both did not change plasma renin activity in our study, suggesting that a more direct action of lisinopril on ACE2-producing cells may be responsible. The precise mechanism by which cells sense angiotensin-II and regulate ACE2 abundance and sub-cellular localization remains to be elucidated.

It is important to note the limitations of this study. All mice used in this study were healthy young adult mice. Hypertension and cardiovascular disease can impact tissue ACE2, and the findings could be different in the setting of cardiovascular disease. While lisinopril and losartan are representative of their drug classes, other ACE inhibitors or ARBs may have different effects on ACE2 abundance. Lastly, while our findings in mice are consistent with the available results in rats, the effects of ACE inhibition and angiotensin receptor blockade on tissue ACE2 levels in humans may be different. A controlled study of tissue ACE2 in humans after initiating an ACE inhibitor, ARB, or combination therapy would be warranted to extend these findings into humans.

ACE2 expression is increased in the lungs of patients with COVID-19 comorbidities^44^, as well as in diabetes ^41,45^ and heart failure.^46^ It is possible that these co-morbidities increase susceptibility to and severity of COVID-19 in part through increased tissue ACE2. In this context, our finding that ACE inhibitor and ARB combination therapy interact to decrease ACE2 gene expression and prevent increases in ACE2 protein levels may offer an avenue to reduce tissue ACE2 in people on ACE inhibitor or ARB monotherapy while still providing protection against cardiovascular or renal disease. While combination therapy of ACE inhibitor with ARB is not widely used, there is precedent for combination therapy for heart failure^47^, renal disease^48^, and among aged individuals^49^. It is important to note that human clinical studies have not identified ACE inhibitors or ARB medications to be risk factors for susceptibility or poor outcome from COVID-19. An analysis of COVID-19 clinical outcomes among people taking ACE inhibitor or ARB monotherapy versus combination therapy could provide valuable observational data about the potential benefits of combination therapy in reducing susceptibility or severity of COVID-19.

## 5. CONCLUSIONS

Lisinopril monotherapy increased ACE2 protein in key tissues affected by SARS-CoV-2, especially the lung and small intestine. In contrast, the combination of lisinopril with losartan prevented the lisinopril-induced increase of tissue ACE2 levels. These results demonstrate that ACE inhibition and angiotensin receptor blockade interact to determine tissue levels of ACE2, the SARS-CoV-2 receptor.

## Supporting information

Graphical abstract

## 6. ACKNOWLEDGEMENTS

We would like to thank Dr. Ian Moore, DVM PhD, Veterinary Pathologist, and Kevin Bock, MS, Histotechnologist, of the Infectious Disease Pathogenesis Section, Comparative Medicine Branch, National Institute of Allergy and Infectious Diseases, for their expertise in performing and interpreting the immunohistochemical staining of ACE2 protein in small intestine, brain, lung, and kidney sections from mice in this project. The graphical abstract was created in part with BioRender (BioRender.com) with the academic publishing rights provided to the National Institute of Allergy and Infectious Diseases.

