## Supplementary material for "Oral Lisinopril Raises Tissue Levels of ACE2, the SARS-CoV-2 Receptor, in Healthy Male and Female Mice": Graphical abstract

### Approach

**Experimental procedure:**

20 Males, 20 Females → Collect tissue and plasma

20 Males, 20 Females → Collect tissue and plasma

Day 0: Start drug

Day 21: Stop drug

**Drug treatment groups:**

- Lisinopril
- Losartan
- Lisinopril + Losartan
- Vehicle control

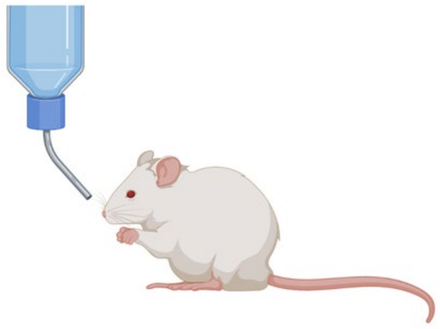

**Tissues evaluated:**

- Small intestine
- Lung
- Kidney
- Brain

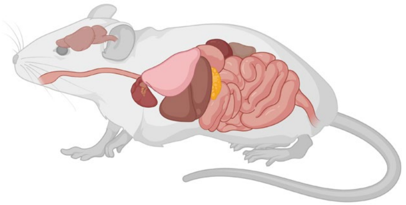

**Primary Endpoint:** $\text{ACE2 Protein Index} = \frac{[\text{Tissue ACE2}]}{[\text{Tissue protein}]}$

### Primary Findings

Oral lisinopril increased ACE2 in tissues.

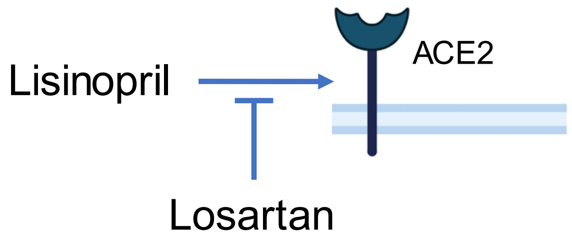


The addition of losartan prevented the increase in tissue ACE2.

### Secondary Findings

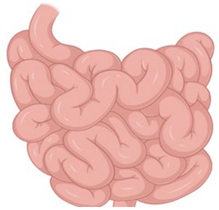

ACE2 index was highest in the small intestine.

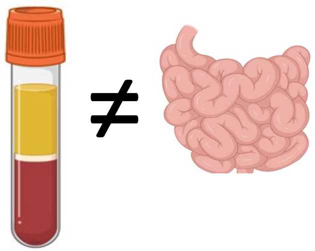

Plasma ACE2 did not correlate with ACE2 in any tissue, indicating it is not a good biomarker for tissue ACE2

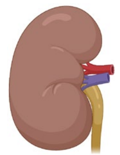

♂ > ♀

In the kidney, ACE2 index was greater in males than in females.

Brooks et al., 2021 *BioRxiv*
