## Appendix A - supplemental materials and methods for "Oral Lisinopril Raises Tissue Levels of ACE2, the SARS-CoV-2 Receptor, in Healthy Male and Female Mice"

**APPENDIX A: DETAILED MATERIALS AND METHODS**

A. **Use of Laboratory Animals**

The protocols used in this study were performed in accordance with the National Institutes of Health guide for the care and use of Laboratory animals (NIH Publications No. 8023, revised 1978). All experiments and protocols using laboratory mice were reviewed and approved by the NIAID Division of Intramural Research Animal Care and Use Committee (DIR ACUC).

Male and Female C57Bl/6J mice (Jackson Laboratories) were utilized in this experiment. All mice were 8 weeks of age at the start of drug treatment.

At the experimental endpoints, mice were euthanized via bilateral thoracotomy while under anesthesia with 4% isoflurane. 1 mL of blood was drawn into an EDTA tube via cardiac puncture of the right ventricle. 25 mL of cold phosphate-buffered saline was administered via transcardial perfusion to remove blood from tissues prior to collection. Small intestine, lung, kidney, and brain were collected and flash-frozen for protein extraction, stored in 10% formalin for histological examination, or stored in RNAlater for gene expression studies. Plasma was separated from whole blood by centrifugation (2000xg, 15 minutes, 4˚C), aliquoted, and frozen until used in ELISA experiments.

B. **Drug treatment with lisinopril and losartan:**

Treatment protocol was approved by NIAID Division of Intramural Research Animal Care and Use Committee (DIR ACUC)

Prior to the main study, a pilot study was conducted to determine the required drug concentrations in drinking water for lisinopril and losartan. For the pilot study, five male and five female eight-week-old C57Bl/6J mice (Jackson Laboratories, strain #000664) were monitored for one week to measure the average daily drinking water intake for male and female mice. This value, combined with the body weight of each mouse, was used to calculate the initial drug concentration needed to achieve a 10mg/kg/day dosage in drinking water. Each week during the main study, water consumption was measured on a per-cage basis, and the body weight of each mouse was recorded. This information was used to adjust the drug concentration weekly to maintain consistent dosing of 10mg/kg/day throughout the course of the study. Body weights (recorded weekly for each animal over the course of the study), and average water consumed by each treatment group during the treatment period, are available in Appendix B: Supplemental Data (Supplemental Fig. 2 for body weight, Supplemental Fig. 3 for water consumption).

The experiment utilized a factorial design: five male and five female mice comprised each drug treatment group (lisinopril, losartan, lisinopril and losartan combined, or vehicle control) at each time point (Day 21 or Day 42). On day 21, forty animals were euthanized for collection of plasma and tissues, while the others transitioned to standard drinking water for an additional twenty-one days to assess whether drug-induced changes in ACE2 resolve after drug cessation.

Lisinopril was obtained from NIH Division of Veterinary Resources Pharmacy as 10mg tablets (Exelan Pharmaceuticals) prepared into suspension in standard drinking water for oral administration. Preparation and storage of oral suspension was conducted in accordance with guidelines from the US Food and Drug Administration (<https://www.accessdata.fda.gov/drugsatfda_docs/label/2009/019777s054lbl.pdf>). New suspensions were formulated weekly to account for changes in mouse body weight and average water consumption.

Losartan was obtained from NIH Division of Veterinary Resources Pharmacy as 25mg tablets (Aurobindo Pharma) prepared into suspension in standard drinking water for oral administration. Preparation and storage of oral suspension was conducted in accordance with guidelines from the US Food and Drug Administration (https://www.accessdata.fda.gov/drugsatfda_docs/label/2013/020386s058lbl.pdf). New suspensions were formulated weekly to account for changes in mouse body weight and average water consumption.

To assess whether treatment with lisinopril, losartan, or combination of both had an impact on the renin-angiotensin system, renin activity was measured in plasma isolated from whole blood from mice at Day 21 and Day 42. Renin activity was measured by a fluorometric Renin Assay Kit (Abcam, Catalog #ab138875). Renin catalytic activity of a fluorescent substrate was measured over time in standards of known renin concentration and in unknown samples by reading fluorescence (excitation at 540 nm, emission at 590 nm) using a microplate reader (Molecular Devices Gemini XPS). The concentration of renin in plasma was extrapolated from a standard curve produced from the activity of the renin standards.

C. **Laboratory equipment and approach for measurement of ACE2 protein index:**

Flash-frozen sections of lung, small intestine, kidney, and brain were homogenized at 4˚C in tubes containing ceramic beads (Precellys CKMix, Catalog # P000918-LYSK0-A) via automated tissue homogenizer (Precellys Cyolys Evolution, Bertin Instruments). Tissue homogenization was performed in lysis buffer (RIPA buffer, working concentration at 1X, Cell Signaling Technology Product #9806S) containing protease/phosphatase inhibitor (1:100, Cell Signaling Technology Product #5872S).

After homogenization, lysed samples were centrifuged (10,000 x g, 20 minutes, 4˚C) to pellet cell debris. Lysate was aliquoted and stored at -80˚C.

Protein concentration for each tissue lysate was determined by Piece BCA Protein Assay (ThermoFisher Scientific, Catalog #23225). Concentration was calculated from absorbance at 562 nm on a plate reader (PerkinElmer VICTOR Nivo Multimode Microplate Reader)

ACE2 tissue abundance was measured by ELISA (Abcam Mouse ACE2 ELISA Kit, product ab213843). Abundance was calculated from absorbance at 450 nm on a plate reader (PerkinElmer VICTOR Nivo Multimode Microplate Reader). To minimize the effects of inter-assay variation, all biospecimens from a given experimental day (21 or 42) were analyzed together on a single ELISA plate and BCA plate.

Tissue ACE2 protein index was calculated by dividing the ELISA-measured concentration by the total protein concentration of each specimen.

ACE2 concentration in plasma (pg/mL) was measured by the same ELISA kit (Abcam Mouse ACE2 ELISA Kit, product ab213843).

D. **Laboratory equipment and approach for measurement of gene expression:**

Tissue sections for RNA extraction were submerged in RNAlater RNA stabilization solution (ThermoFisher Scientific, catalog #AM7021M) and then stored at -80˚C.

RNA extraction was performed using the RNeasy Mini Kit (Qiagen, Catalog # 74106) and RNA concentration and quality was measured by DeNovix (DeNovix DS-11 FX+). RNA was then converted to cDNA (SuperScript™ IV VILO™ Master Mix with ezDNase™ Enzyme, ThermoFisher Scientific Catalog #11766050) and diluted to a standard concentration of 25 ng/μL.

Gene expression of *Ace2* and *Gapdh* was measured by droplet digital polymerase chain reaction (ddPCR). A total of 1 ng cDNA per sample was loaded into a reaction well with ddPCR Supermix for Probes (No dUTP) (Bio-Rad, Catalog #1863024) and commercially available ddPCR gene expression primer/probe assays for murine *Ace2* (Bio-Rad, unique assay ID: dMmuCPE5123530) and murine *Gapdh* (Bio-Rad, unique assay ID: dMmuCPE5195283). Droplets for ddPCR were created by automated droplet generator (Bio-Rad QX200 AutoDG); amplification was performed with T100 Thermal Cycler (Bio-Rad); and ddPCR droplet reading was performed on automated droplet reader (Bio-Rad Rad QX200 Droplet Reader). ddPCR gene expression results were analyzed by QuantaSoft Analysis Pro (Bio-Rad).

E. **Laboratory equipment and approach for tissue histology and immunohistochemistry (IHC)**

Mouse tissue samples were fixed in 10% neutral buffered formalin and then processed via Leica ASP6025 tissue processor (Leica Biosystems). Tissues were embedded in paraffin, and sectioned by microtome at 5 μm thickness.

Tissue sections were stained with hematoxylin and eosin (H&E) for routine histopathology.

For immunohistochemical (IHC) evaluation, sections were labeled with rabbit polyclonal anti-ACE2 antibody at 1:1000 (Sino Biological, catalog #10108-T24). Chromogenic staining was carried out on the Bond RX (Leica Biosystems) platform according to manufacturer-supplied protocols. Formalin-fixed paraffin-embedded (FFPE) sections were deparaffinized and rehydrated. Heat-induced epitope retrieval (HIER) was performed using the citrate-based Epitope Retrieval Solution 1 (Leica Biosystems AR9961), pH 6.0, heated to 100˚C for 20 minutes. The specimen was then incubated with hydrogen peroxide to quench endogenous peroxidase activity prior to applying the primary antibody for 15 minutes. Detection with DAB chromogen was completed using the Bond Polymer Refine Detection kit (Leica Biosystems DS9800). Slides were finally cleared through gradient alcohol and xylene washes prior to mounting and coverslipping. Sections were examined by light microscopy using an Olympus BX51 microscope and photomicrographs were taken using an Olympus DP73 camera.
