## Appendix B - supplemental data tables and figures for "Oral Lisinopril Raises Tissue Levels of ACE2, the SARS-CoV-2 Receptor, in Healthy Male and Female Mice"

**APPENDIX B: SUPPLEMENTAL TABLES AND FIGURES**

| **Day** | **Tissue** | **Treatment** | **n** | **Mean** | **SD** |
| --- | --- | --- | --- | --- | --- |
| 21 | Brain | Vehicle | 10 | 6.50 | 1.08 |
| 21 | Brain | Lisinopril | 10 | 7.20 | 0.73 |
| 21 | Brain | Losartan | 10 | 6.43 | 0.85 |
| 21 | Brain | Combination | 10 | 6.36 | 1.12 |
| 21 | Kidney | Vehicle | 10 | 71.2 | 24.4 |
| 21 | Kidney | Lisinopril | 10 | 89.9 | 23.5 |
| 21 | Kidney | Losartan | 10 | 75.2 | 21.7 |
| 21 | Kidney | Combination | 10 | 79.6 | 25.8 |
| 21 | Lung | Vehicle | 10 | 7.84 | 2.48 |
| 21 | Lung | Lisinopril | 10 | 11.4 | 4.12 |
| 21 | Lung | Losartan | 10 | 9.75 | 2.96 |
| 21 | Lung | Combination | 10 | 7.40 | 1.43 |
| 21 | Sm. Intestine | Vehicle | 10 | 187 | 92.7 |
| 21 | Sm. Intestine | Lisinopril | 10 | 375 | 128 |
| 21 | Sm. Intestine | Losartan | 10 | 288 | 111 |
| 21 | Sm. Intestine | Combination | 10 | 234 | 109 |
| 42 | Brain | Vehicle | 10 | 4.94 | 0.811 |
| 42 | Brain | Lisinopril | 10 | 6.12 | 0.525 |
| 42 | Brain | Losartan | 10 | 5.99 | 1.27 |
| 42 | Brain | Combination | 10 | 5.95 | 0.938 |
| 42 | Kidney | Vehicle | 10 | 47.0 | 14.5 |
| 42 | Kidney | Lisinopril | 9 | 63.7 | 21.5 |
| 42 | Kidney | Losartan | 10 | 63.9 | 18.9 |
| 42 | Kidney | Combination | 10 | 63.9 | 12.1 |
| 42 | Lung | Vehicle | 10 | 6.63 | 1.21 |
| 42 | Lung | Lisinopril | 9 | 8.63 | 2.71 |
| 42 | Lung | Losartan | 10 | 9.35 | 2.11 |
| 42 | Lung | Combination | 10 | 7.91 | 2.85 |
| 42 | Sm. Intestine | Vehicle | 10 | 221 | 70.7 |
| 42 | Sm. Intestine | Lisinopril | 9 | 251 | 87.9 |
| 42 | Sm. Intestine | Losartan | 10 | 237 | 107 |
| 42 | Sm. Intestine | Combination | 10 | 230 | 120 |

**Supplemental Table 1: Tissue ACE2 protein index.** ACE2 protein index was determined in tissues from male and female mice in each treatment group. Protein index was calculated as the ratio of ACE2 protein to total protein. Displayed values have been multiplied by 10^6^. At Day 42, one female mouse was removed from the lisinopril group due to a congenital defect.

| **Day** | **Tissue** | **Treatment** | **n** | **Mean** | **SD** |
| --- | --- | --- | --- | --- | --- |
| 21 | Brain | Vehicle | 10 | 0.000558 | 0.0000917 |
| 21 | Brain | Lisinopril | 8 | 0.000477 | 0.0000788 |
| 21 | Brain | Losartan | 10 | 0.000499 | 0.000134 |
| 21 | Brain | Combination | 10 | 0.000459 | 0.000128 |
| 21 | Kidney | Vehicle | 10 | 0.0134 | 0.00157 |
| 21 | Kidney | Lisinopril | 9 | 0.0126 | 0.00152 |
| 21 | Kidney | Losartan | 10 | 0.0134 | 0.00147 |
| 21 | Kidney | Combination | 10 | 0.0132 | 0.00112 |
| 21 | Lung | Vehicle | 10 | 0.0162 | 0.00423 |
| 21 | Lung | Lisinopril | 10 | 0.0167 | 0.00307 |
| 21 | Lung | Losartan | 9 | 0.0172 | 0.00446 |
| 21 | Lung | Combination | 10 | 0.0185 | 0.00397 |
| 21 | Sm. Intestine | Vehicle | 10 | 0.400 | 0.131 |
| 21 | Sm. Intestine | Lisinopril | 10 | 0.305 | 0.0964 |
| 21 | Sm. Intestine | Losartan | 9 | 0.286 | 0.0478 |
| 21 | Sm. Intestine | Combination | 10 | 0.181 | 0.116 |

**Supplemental Table 2: *Ace2*** **Gene Expression.** *Ace2* gene expression was measured as the ratio of *Ace2/Gadph* in male and female mice from each treatment group at day 21. Three outliers were removed before analysis.

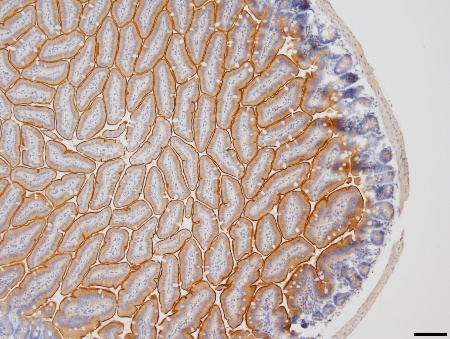

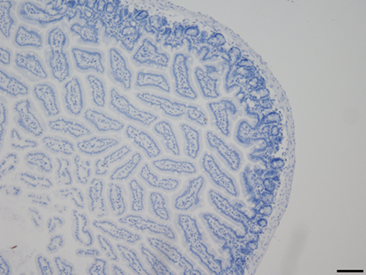

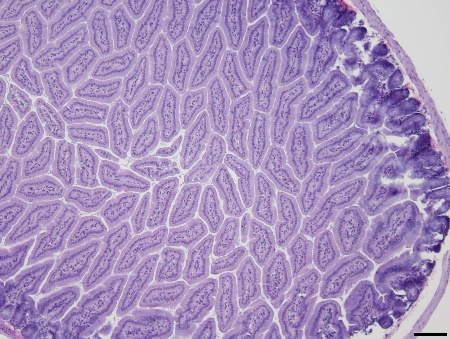

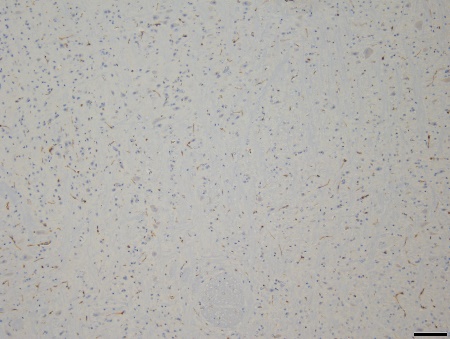

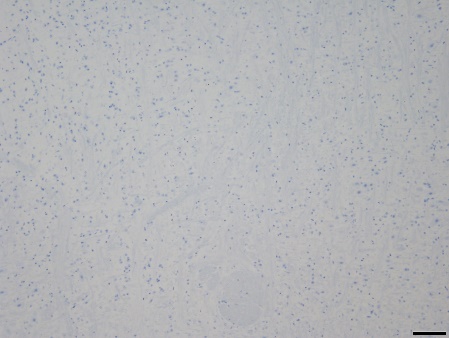

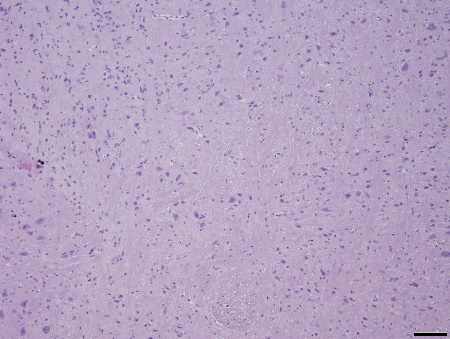

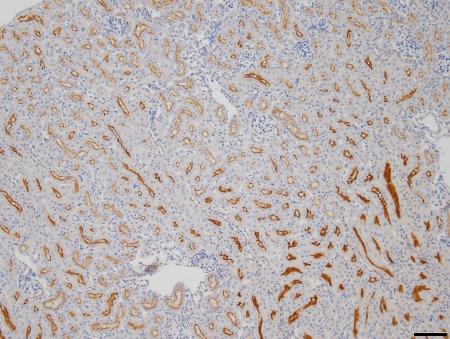

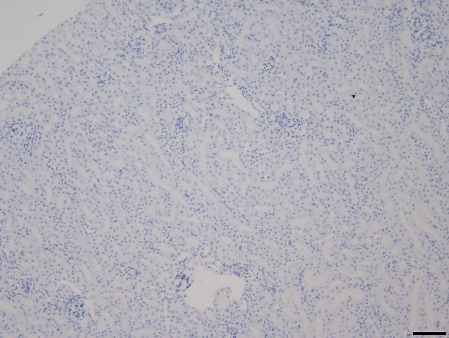

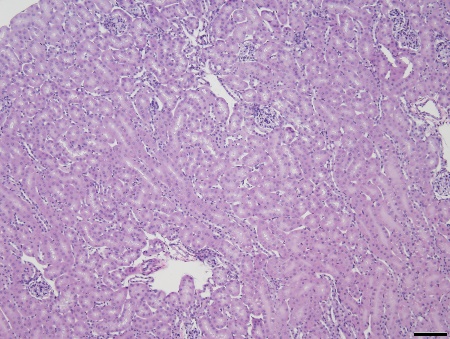

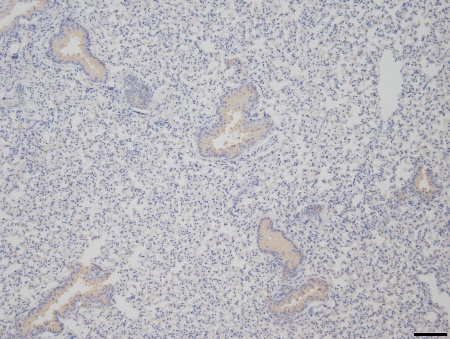

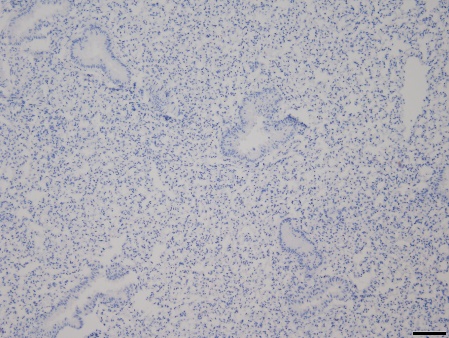

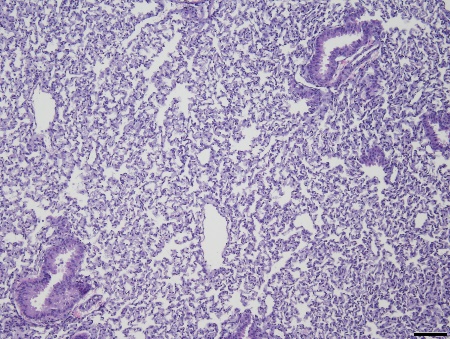

**ACE2**

**Negative Control**

**Hematoxylin & Eosin**

**Small Intestine**

**Kidney**

**Lung**

**Brain**

**Supplemental Figure 1**: **Immunohistochemical staining of ACE2 in four tissues.** ACE2 immunohistochemistry of small intestine, kidney, lung, and brain from a vehicle-treated male mouse at Day 21. Scale bar = 100 µm. The primary antibody was omitted from the negative control sections. Hematoxylin and eosin-stained sections are provided for comparison.

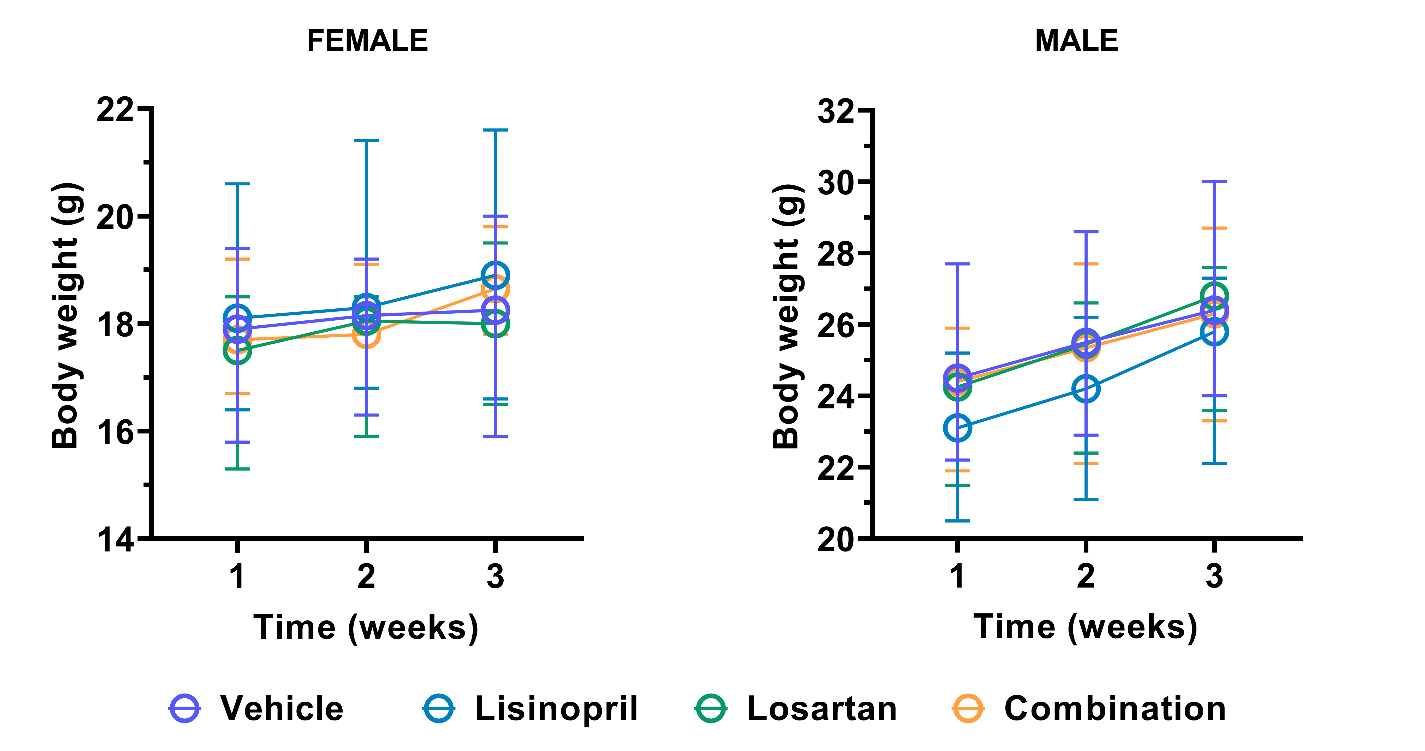

**B**

**A**

**Supplemental Figure 2: Mouse body weight increased over time.** To determine if treatment affected body weight, the mice were weighed once per week during the 21-day treatment period. Each treatment group had 10 mice; data are presented as median and range. The effect of time and treatment on body weight was determined by two-way ANOVA with repeated measures. In female mice, body weight increased over time (p < 0.0001) but was not affected by treatment (p = 0.360). In male mice, body weight increased over time (p < 0.0001) but was not affected by treatment (p = 0.107). There was no interaction between time and treatment in females (p = 0.673) or males (p = 0.084).

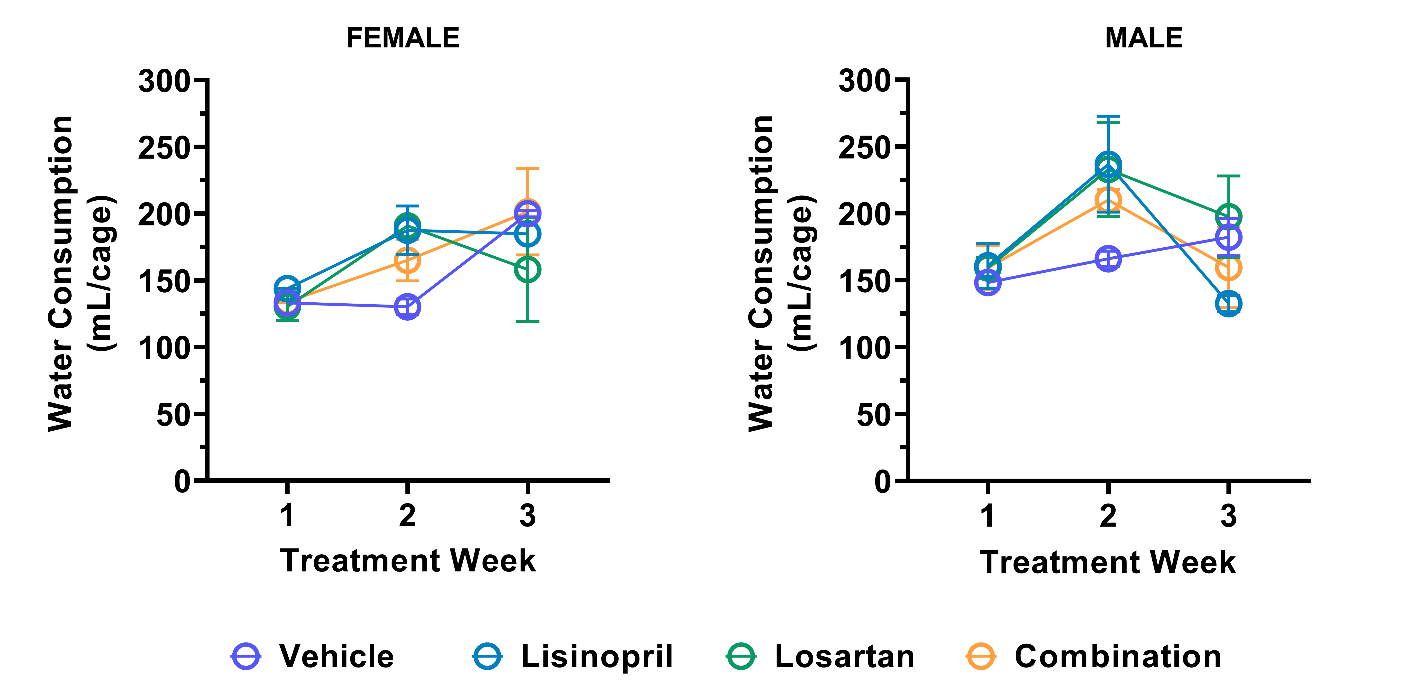

**B**

**A**

**Supplemental Figure 3: Water consumption by treatment group.** To measure if treatment had an impact on water consumption, the weekly intake of water was tracked on a per-cage basis for female (A) and male (B) mice over the 21-day treatment period. Each treatment group had 10 mice (5 mice in each cage, two cages per treatment group). Data are presented as median and range. The effect of time and treatment on water consumption was determined by two-way ANOVA with repeated measures. In female mice, water consumption changed over time (p = 0.015) but was not affected by treatment (p = 0.885). In male mice, water consumption changed over time (p = 0.050) but was not affected by treatment (p = 0.436). There was no interaction between time and treatment and in females (p = 0.116) or males (p = 0.279).

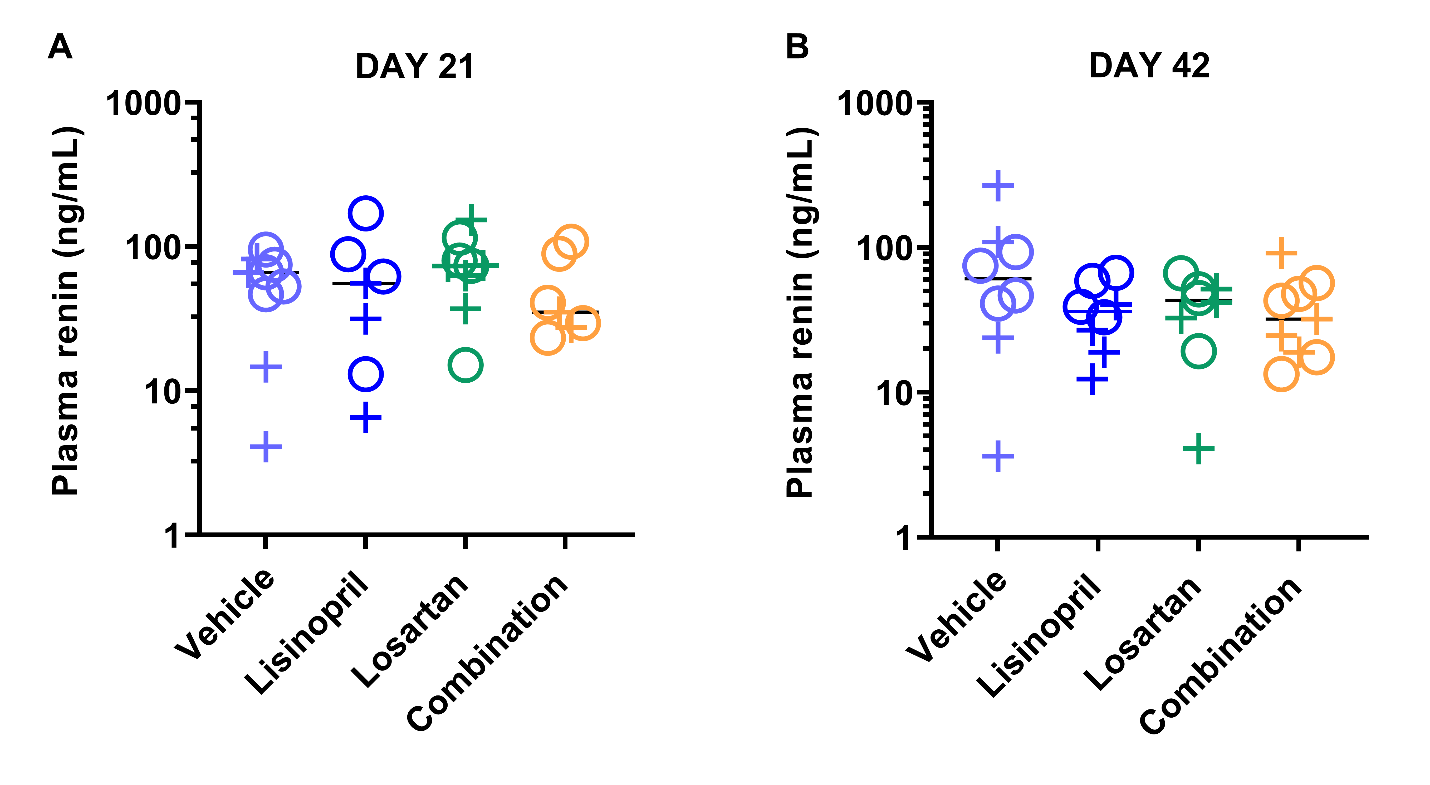

**Supplemental Figure 4: Plasma Renin Activity.** To assess whether treatment had an impact on activation of the renin-angiotensin system, plasma samples from mice at day 21 (A) and day 42 (B) were assayed for renin activity. The activity of each plasma sample was extrapolated against catalytic activity from renin standards of known concentration and presented as plasmin renin ng/mL equivalent. Groups included 5 males and 5 females except day 42 female losartan in which n=4. The effects of treatment and sex was assessed by two-way ANOVA performed separately on day 21 and day 42 data. At day 21, neither treatment (p = 0.515) nor sex (p = 0.120) affected plasma renin activity. At day 42, neither treatment (p = 0.166) nor sex (p = 0.935) affected plasma renin activity.

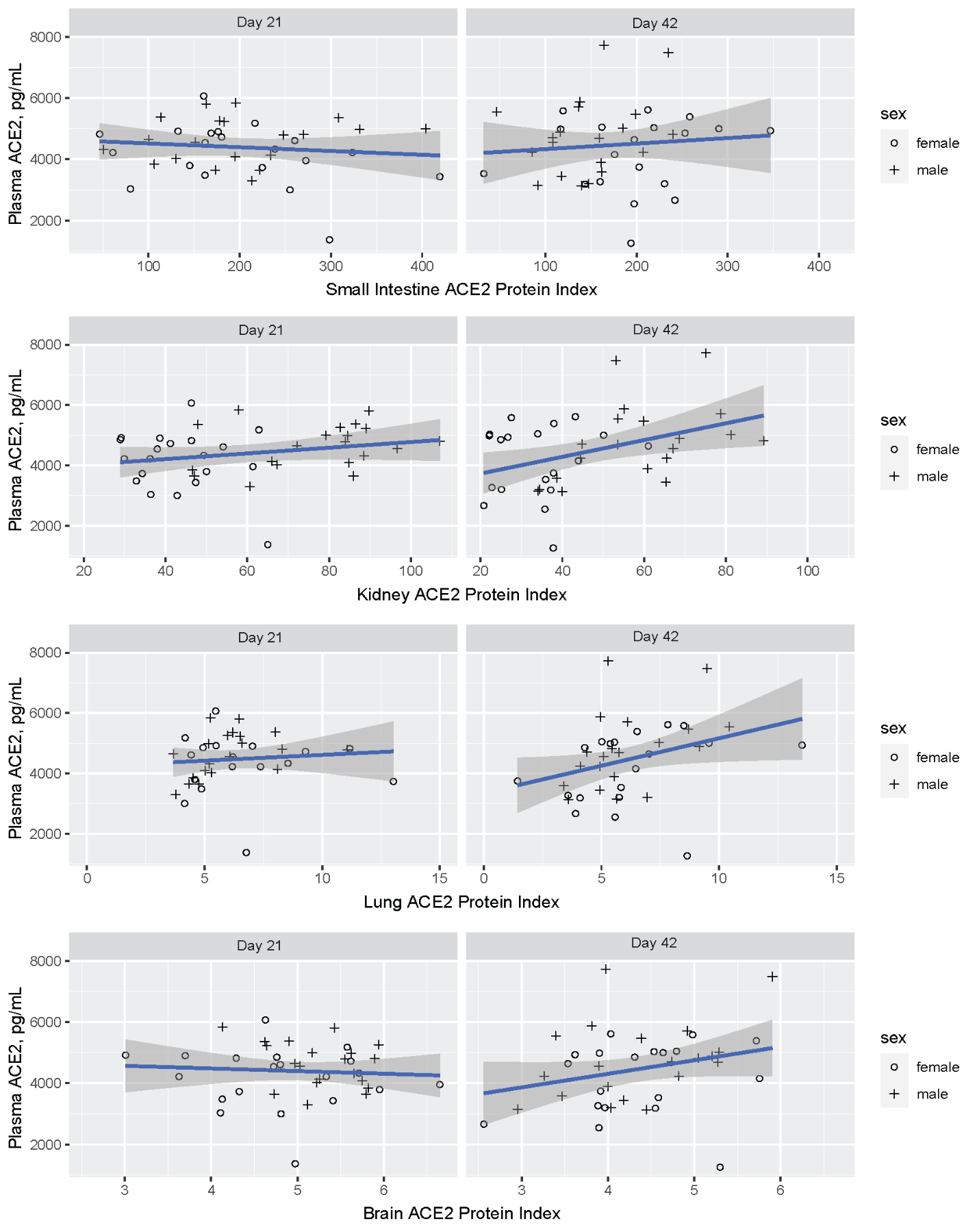
**Supplemental Figure 5: Plasma ACE2 concentration is not correlated with tissue ACE2 protein index in any tissue.** Plasma ACE2 concentration (pg/mL) was determined by ELISA and plotted against the ACE2 protein index for each tissue (small intestine, kidney, lung, brain). Data from female (o) and male (+) mice are shown together. Day 21 and day 42 results are plotted separately. The relationships between plasma ACE2 and tissue ACE2 protein index in small intestine, lung, kidney, and brain were analyzed by multivariable linear regression. Separate models were run for each tissue with terms included for sex, treatment group, and experimental day. Plasma ACE2 was not associated with tissue ACE2 index in any tissue (small intestine p = 0.95; kidney p = 0.26; lung p = 0.90; brain p = 0.62). ACE2 protein indices were multiplied by 10^6^ for display purposes.
